# Reproducible and Transparent Research Practices in Published Neurology Research

**DOI:** 10.1101/763730

**Authors:** Shelby Rauh, Trevor Torgerson, Austin L. Johnson, Jonathan Pollard, Daniel Tritz, Matt Vassar

**Author notes:** Corresponding Author: Shelby Rauh 1111 W 17th St. Tulsa, OK 74137. **Data Availability Statement**: All protocols, materials, and raw data are available online via bioRxiv (BIOARKIV/2019/763730).

## Abstract

**Background:** The objective of this study was to evaluate the nature and extent of reproducible and transparent research practices in neurology research.

**Methods:** The NLM catalog was used to identify MEDLINE-indexed neurology journals. A PubMed search of these journals was conducted to retrieve publications over a 5-year period from 2014 to 2018. A random sample of publications was extracted. Two authors conducted data extraction in a blinded, duplicate fashion using a pilot-tested Google form. This form prompted data extractors to determine whether publications provided access to items such as study materials, raw data, analysis scripts, and protocols. In addition, we determined if the publication was included in a replication study or systematic review, was preregistered, had a conflict of interest declaration, specified funding sources, and was open access.

**Results:** Our search identified 223,932 publications meeting the inclusion criteria, from which 300 were randomly sampled. Only 290 articles were accessible, yielding 202 publications with empirical data for analysis. Our results indicate that 8.99% provided access to materials, 9.41% provided access to raw data, 0.50% provided access to the analysis scripts, 0.99% linked the protocol, and 3.47% were preregistered. A third of sampled publications lacked funding or conflict of interest statements. No publications from our sample were included in replication studies, but a fifth were cited in a systematic review or meta-analysis.

**Conclusions:** Current research in the field of neurology does not consistently provide information needed for reproducibility. The implications of poor research reporting can both affect patient care and increase research waste. Collaborative intervention by authors, peer reviewers, journals, and funding sources is needed to mitigate this problem.

## Background

Scientific advancement is hampered by potential research flaws, such as the lack of replication; poor reporting; selective reporting bias; low statistical power; and inadequate access to materials, protocols, analysis scripts, and experimental data.[1–3] These factors may undermine the rigor and reproducibility of published research. Substantial evidence suggests that a large proportion of scientific evidence may be false, unreliable, or irreproducible.[4–8] Estimates of irreproducible research range from 50% to 90% in preclinical sciences[9] and substantiated in a recent survey of scientists. Prior survey studies reported that roughly 70% of scientists were unable to replicate another scientist’s experiment, and 90% agreed that scientific research is currently experiencing a “reproducibility crisis.”[7]

Reproducibility is vital for scientific advancement as it aids in enhancing the credibility of novel scientific discoveries and mitigates erroneous findings. One review discussed potential pitfalls in fMRI reproducibility, such as scanner settings, consistency of cognitive tasks, and analysis methods.[10] Boekel et al. replicated five fMRI studies measuring a total of 17 structural brain-behavior correlations. After reanalysis, only one of the 17 was successfully replicated.[11] Thus, practices related to transparency and reproducibility can be improved within fMRI and other neurology research.

Adopting open science in neurology would help mitigate irreproducible research, such as the studies on brain-behavior correlation. Open science practices – such as data sharing, open access articles, sharing protocols and methods, and study preregistration – promote transparency and reproducibility.[12] For example, preregistering a study helps guard against selective outcome reporting.[13] Selective outcome reporting occurs when discrepancies exist between outcome measures prespecified in trial registries or research protocols and the outcomes listed in the published report.[14] In neurology, an audit of randomized clinical trials published in neurology journals found 180 outcome inconsistencies across 180 trials, with most inconsistencies favoring changes in accordance with statistically significant results. Additionally, only 55% of neurology trials were prospectively registered[15], providing indications that neurology researchers are not adhering to transparency and reproducibility practices early in research planning. Reproducible research and open science practices are widely endorsed by a large proportion of authors. Despite this support, evidence suggests that authors infrequently implement them.[16–18]

Given the recent attention to the reproducibility crisis in science, further investigation is warranted to ensure the existence of reproducible and transparent research in the field of neurology. Here, we examine key transparency- and reproducibility-related research practices in the published neurology literature. Our findings from this investigation may serve as a baseline to measure future progress regarding transparency and reproducibility-related practices.

## Methods

This observational, cross-sectional study used the methodology proposed by Hardwicke et. al.[3], with modifications. We reported this study in accordance with the guidelines for meta-epidemiological methodology research[19] and, when pertinent, the Preferred Reporting Items for Systematic Reviews and Meta-Analyses (PRISMA).[20] Our study did not use any human subjects or patient data and, as such, was not required to be approved by an institutional review board prior to initiation. We have used The Open Science Framework to host our protocol, materials, training video, and study data in a publically available database (https://osf.io/n4yh5/).

### Journal and Publication Selection

On June 25, 2019, one investigator (D.T.) searched the National Library of Medicine (NLM) catalog for all journals using the subject terms tag “Neurology[ST].” The inclusion criteria required that all journals publish English, full-text manuscripts and be indexed in the MEDLINE database. The final list of included journals was created by extracting the electronic international standard serial number (ISSN) or the linking ISSN, if necessary. PubMed was searched with the list of journal ISSNs on June 25, 2019 to identify all publications. We then limited our publication sample to those between January 1, 2014 and December 31, 2018. Three hundred publications within the time period were randomly sampled for data extraction. The rest were available, but not needed (https://osf.io/wvkgc/).

### Extraction Training

Prior to data extraction, two investigators (S.R. and J.P.) completed in-person training designed and led by another investigator (D.T.). The training sessions included reviewing the protocol, study design, data extraction form, and likely locations of necessary information within example publications. The two authors being trained received two sample publications to extract data from. This example data extraction was performed in the same duplicate and blinded fashion used for data acquisition for this study. The two investigators then met to reconcile any discrepancies. After the two sample publications were completed, investigators extracted data and reconciled differences from the first 10 of the included 300 neurology publications. This process insured interrater reliability prior to analyzing the remaining 290 publications. A final reconciliation meeting was conducted, with a third investigator (D.T.) available for disputes but not needed.

### Data Extraction

After completing training, the same two investigators extracted data from the included list of randomly sampled publications between June 3, 2019 and June 10, 2019 using a pilot-tested Google form. This Google form was based on the one used by Hardwicke et al., but including modifications.[3] We specified the 5-year impact factor and that for the most recent year as opposed to the impact factor of a specific year. The available types of study designs were expanded to include case series, cohort studies, secondary analyses, chart reviews, and cross-sectional analyses. Last, we specified funding sources, such as hospital, private/industry, non-profit, university, or mixed, instead of restricting the criteria to public or private.

### Assessment of Reproducibility and Transparency Characteristics

This study used the methodology by Hardwicke et al.[3] for analyses of transparency and reproducibility of research, with modifications. Full publications were examined for funding disclosures, conflicts of interest, available materials, data, protocols, and analysis scripts. Publications were coded to fit two criteria: those with and those without empirical data. Publications without empirical data (e.g., editorials, reviews, news, simulations, or commentaries without reanalysis) were analyzed for conflict of interest statements, open access, and funding. Given that protocols, data sets, and reproducibility were not relevant, these were omitted. Case studies and case series were listed as empirical studies; however, questions pertaining to the availability of materials, data, protocol, and registration were excluded due to previous study recommendations.[18] Data extraction criteria for each study design is outlined in Table 1.

**Table 1:** Reproducibility related characteristics. Variable numbers (N) are dependent upon study design. Full detailed protocol pertaining to our measured variables is available online (https://osf.io/x24n3/)

| *Indicators of Reproducibility Included in Present Study* |  | *Significance of measure variable for transparency and reproducibility.* |
| --- | --- | --- |
| **Publications** |  |  |
| All<br>(N=300) | Publication accessibility (Is the publication open access to the general public or accessible through a paywall?) | The general public's ability to access scientific research may increase transparency of results and improve the ability for others to critically assess studies, potentially resulting in more replication studies |
| **Funding** |  |  |
| Included studies<br>(N=290) | Funding statement (Does the publication state their funding sources?) | Explicitly providing source of funding may help mitigate bias and potential conflicts of interest |
| **Conflict of Interest** |  |  |
| Included studies<br>(N=290) | Conflict of interest statement (Does the publication state whether or not the authors had a conflict of interest?) | Explicitly providing conflicts of interest may allow for full disclosure of factors that may promote bias in the study design or outcomes |
| **Publication Citations** |  |  |
| Empirical studies† | Citations by a systematic review/meta-analysis (Has the publication been | Systematic reviews and meta-analyses evaluate and compare existing literature to assess for |
| (N=205) | cited by any type of data synthesis publication, and if so, was it explicitly excluded?) | patterns, strengths, and weaknesses of studies regarding a particular field or topic |
| Analysis Scripts |  |  |
| Empirical studies‡<br>(N=202) | Availability statement (Does the publication state whether or not the analysis scripts are available?) | Providing access to the analysis script helps improve credibility by providing the replicators the opportunity to analyze raw data with the same analysis procedure |
|  | Method of availability (Ex: Are the analysis scripts available upon request or in a supplement?) |  |
|  | Accessibility (Can you view, download, or otherwise access the analysis scripts?) |  |
| Materials |  |  |
| Empirical studies¶<br>(N=189) | Availability statement (Does the publication state whether or not the materials are available?) | Providing the materials list allows replicators to reproduce study using the same materials, promoting |
|  | Method of availability (Ex: Are the materials available upon request or in a supplement?) |  |
|  | Accessibility (Can you view, download, or otherwise access the materials?) |  |

| Pre-registration |  |  |
| --- | --- | --- |
| Empirical studies‡<br>(N=202) | Availability statement (Does the publication state whether or not it was pre-registered?) | Pre-registering studies may help mitigate potential bias and increase the overall validity and reliability of a study |
|  | Method of availability (Where was the publication pre-registered?) |  |
|  | Accessibility (Can you view or otherwise access the registration?) |  |
|  | Components (What components of the publication were pre-registered?) |  |
| Protocols |  |  |
| Empirical studies‡<br>(N=202) | Availability statement (Does the publication state whether or not a protocol is available?) | Providing replicators access to protocols allows the for more accurate replication of the study, promoting credibility |
|  | Components (What components are available in the protocol?) |  |
| Raw Data |  |  |
| Empirical studies‡<br>(N=202) | Availability statement (Does the publication state whether or not the raw data are available?) | Providing replicators with access to raw data can help reduce potential bias and increase validity and reliability |
|  | Method of availability (Ex: Are the raw data available upon request or in a supplement?) |  |
|  | Accessibility (Can you view, download, or otherwise access the raw data?) |  |
|  | Components (Are all the necessary raw data to reproduce the study available?) |  |
|  | Clarity (Are the raw data documented clearly?) |  |
| † 'Empirical studies' are publications that include empirical data such as: clinical trial, cohort, case series, case reports, case-control, secondary analysis, chart review, commentaries (with data analysis), laboratory, surveys, and cross-sectional designs.<br>‡ Empirical studies determined to be case reports or case series were excluded in regard to reproducibility related questions (materials, data, protocol, and registration were excluded) as recommended by Wallach et al.<br>¶ Empirical studies determined to be either case reports, case series, commentaries with analysis, meta-analysis or systematic review were excluded as they did not provide materials to fit the category. |  |  |

### Publication Citations Included in Research Synthesis and Replication

For both empirical and nonempirical studies, we measured the impact factor of each journal by searching for the publication title on the Web of Science (https://webofknowledge.com). For empirical studies, we used the Web of Science to determine whether our sample of studies were cited in either a meta-analysis, systematic review, or a replication study. The Web of Science provided access to studies that cited the queried publication and provided the title, abstract, and link to the full-text article. This permitted evaluation of the inclusion of the queried article in data synthesis. Extraction was performed by both investigators in a duplicate, blinded fashion.

### Assessment of Open Access

Important core components of publications necessary for reproducibility are only available within the full text of a manuscript. To determine the public’s access to each publication’s full text, we systematically searched the Open Access Button (https://openaccessbutton.org), Google, and PubMed. First, we searched the title and DOI using the Open Access Button to determine if the publication was available for public access. If this search returned no results or had an error, then we searched the publication title on Google or PubMed and reviewed the journal website to determine if the publication was available without a paywall.

### Statistical Analysis

Microsoft Excel was used to report statistics for each category of our analysis. In particular, we used Excel functions to calculate our study characteristics, results, and 95% confidence intervals.

## Results

### Journal and Publication Selection

After searching the National Library of Medicine catalog, 490 neurology journals were eligible for analysis. After screening for inclusion criteria, 299 journals remained for analysis, yielding 223,932 publications. Of the 223,932 publications, we randomly sampled 300 (https://osf.io/qfy7u/). Ten publications were inaccessible, which left 290 publications for analysis. Of the 290 eligible publications, 218 provided analyzable empirical data, and 72 articles were excluded because they did not contain characteristics measurable for reproducibility. Of the 218 publications eligible for analysis, an additional 16 case studies and case series were excluded, as they are irreproducible. Our final analysis was based on 202 publications with measurable reproducibility characteristics (Figure 1 and Table 1).

**Figure 1:**
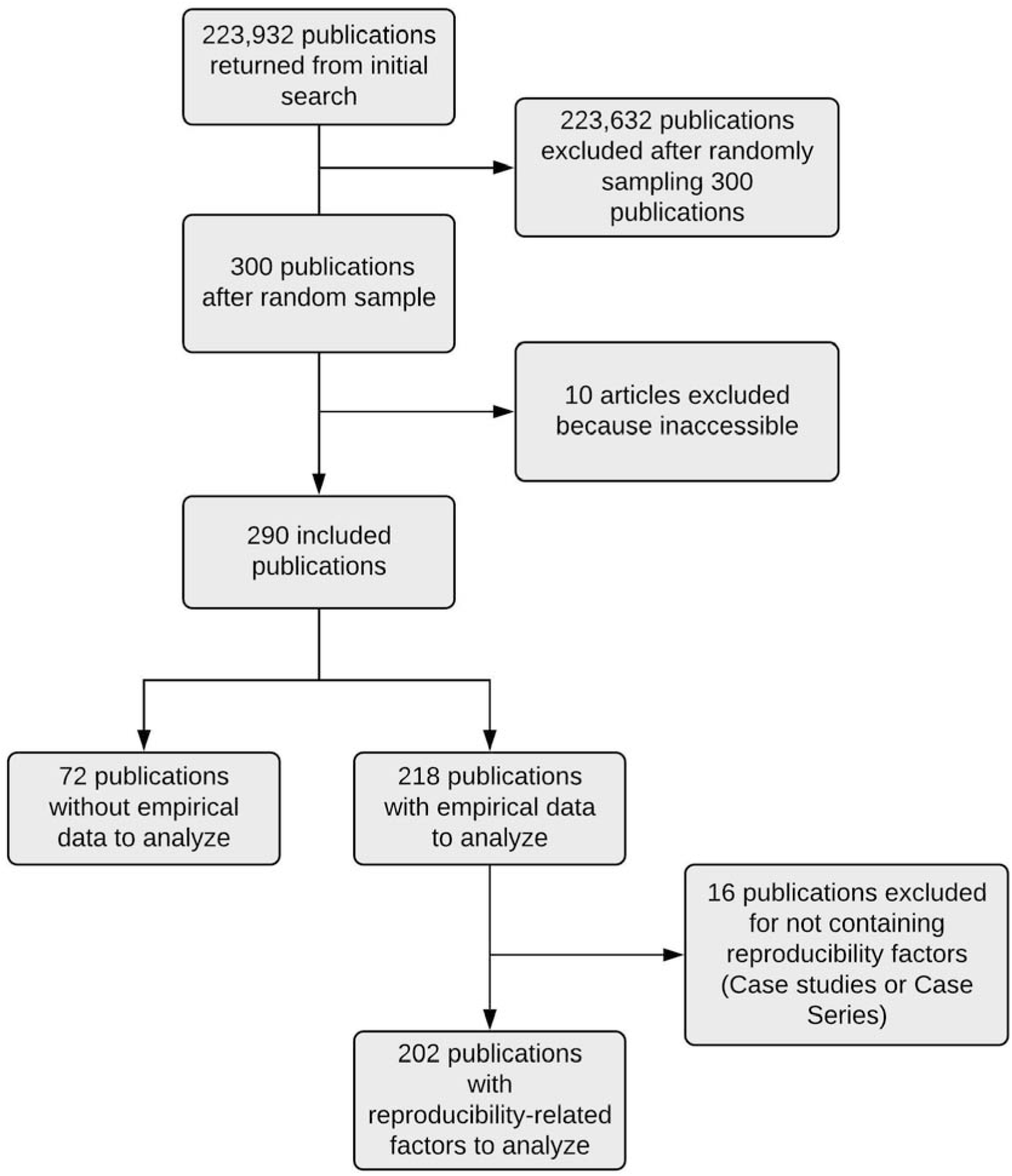
Flow Diagram of Included and Excluded Studies for the Reproducibility Analysis

### Sample Characteristics

Of the eligible publications, the median 5-year impact factor was 3.555 (Interquartile range (IQR): 2.421-4.745), although 20 publications had inaccessible impact factors. The United States was the location of most of the primary authors (30.69%, 89/290) and the country of most publications (56.55%, 164/290). Of the 290 publications that were accessible, 33.10% did not report a funding source (96/290), and 27.93% reported funding from mixed sources (81/290; Table 2).

**Table 2:** Characteristics of Included Publications

| **Table 2: Characteristics of Included Publications** |  |  |  |
| --- | --- | --- | --- |
| **Characteristics** |  | **Variables** |  |
|  |  | **N (%)** | **95% CI** |
| **Funding**<br>**N=290** | University | 11 (3.79) | 1.63-5.95% |
|  | Hospital | 1 (0.34) | 0-1.01% |
|  | Public | 54 (18.62) | 14.22-23.03% |
|  | Private/Industry | 15 (5.17) | 2.67-7.68% |
|  | Non-Profit | 11 (3.79) | 1.63-5.95% |
|  | Mixed | 81 (27.93) | 28.89-30.53% |
|  | No Statement Listed | 96 (33.01) | 27.78-38.43% |
|  | No Funding Received | 21 (7.24) | 4.31-10.17% |
| **Type of Study**<br>**N=290** | No Empirical Data | 72 (24.83) | 19.94-29.72% |
|  | Meta-Analysis | 12 (4.14) | 1.88-6.39% |
|  | Commentary with Analysis | 1 (0.34) | 0-1.01% |
|  | Cost-Effectiveness | 1 (0.34) | 0-1.01% |
|  | Clinical Trial | 26 (8.97) | 5.73-12.20% |
|  | Case Study | 9 (3.10) | 1.14-5.07% |
|  | Case Series | 7 (2.41) | 0.68-4.15% |
|  | Cohort | 43 (14.83) | 10.81-18.85% |
|  | Chart Review | 3 (1.03) | 0-2.18% |
|  | Case Control | 9 (3.10) | 1.14-5.07% |
|  | Survey | 5 (1.72) | 0.25-3.20% |
|  | Cross-Sectional | 34 (11.72) | 8.08-15.36% |
|  | Secondary Analysis | 2 (0.69) | 0-1.63% |
|  | Laboratory | 65 (22.41) | 17.69-27.13% |
|  | Multiple Study Types | 1 (0.34) | 0-1.01% |
| **5 Year Impact**<br>**Factor**<br>**N=278** | Median | 3.555 | - |
|  | 1st Quartile | 2.421 | - |
|  | 3rd Quartile | 4.745 | - |
|  | Interquartile Range | 2.421-4.745 | - |

Of the randomly sampled 300 publications that were findable, 61.38% were not accessible to the public without a paywall (178/290), and only 40.34% were available to the public via the Open Access Button (117/290). Approximately half of analyzed publications stated that they did not have any conflicts of interest (53.10%, 154/290), and 33.10% did not report whether or not conflicts of interest existed (96/290). Humans were the focus of 51.72% of the analyzed publications (150/290). Additional sample characteristics are viewable in Supplemental Tables 1, 2, and 3.

### Reproducibility-Related Characteristics

Among the 202 publications with empirical data that were analyzed, a mere 3.47% provided preregistration statements or claimed to be preregistered (7/202). Of the 202 publications, just 0.99% provided access to the protocol (2/202). Interestingly, only 8.99% provided access to the materials list (17/189), 9.41% provided access to the raw data (19/202), and just a single article provided the analysis script (0.50%, 1/202). Not a single publication claimed to be a replication study. Additional characteristics are viewable in Supplemental Tables 1, 2, and 3.

## Discussion

Our analysis demonstrates inadequate reproducibility practices within neurology and neuroscience research. We found that few publications contained data or materials availability statements and even fewer contained a preregistration statement, made the protocol available, or included an analysis script. Our overall finding – that a majority of neurology publications lack the information necessary to be reproduced and transparent – is comparable to findings in the social and preclinical sciences.[3, 5, 21–24] Here, we present a discussion on prominent reproducibility and transparency indicators that were lacking in our study while presenting recommendations and practices to help improve neurology research.

First, data and materials availability is essential for reproducing research. Without source data, corroborating the results is nearly impossible. Without a detailed description of materials, conducting the experiment becomes a guessing game. Less than 10% of publications in our sample reported either a data or a materials availability statement. Efforts toward data sharing in neurological research originated with brain mapping and neuroimaging, but has spread to other areas within the specialty to improve reproducibility, transparency, and data aggregation.[25] Although data sharing poses challenges, steps have been taken in fMRI studies.[26, 27] fMRI data are complex and cumbersome to handle, but can be managed with software, such as Automatic Analysis[28], C-BRAIN[29], and NeuroImaging Analysis Kit.[30] Furthermore, these data can be hosted on online repositories, such as The National Institute of Mental Health Data Archive[31], Figshare[32], and other National Institutes of Health repositories.[33] Although researchers may take these steps voluntarily, journals – the final arbiters of research publications – can require such practices. Our study found that less than half of the sampled journals had a data availability policies, with approximately 20% of articles from these journals reporting source data.[34] Another study in *PLOS ONE* found that only 20% of nearly 50,000 publications included a data sharing statement and found that once a data sharing policy was enacted, open access to raw data increased.[35] Based on this evidence, journals and funders should consider implementing and enforcing data sharing policies that, at minimum, require a statement detailing whether data are available and where data are located. For example, the journal *Neurology* has endorsed the International Committee of Medical Journal Editors policy of requiring a data sharing statement and encourages open access.[36–38] If other neurology journals follow suit, an environment of transparency and reproducibility may be established.

Second, preregistration practices were uncommon among neurology researchers. Preregistration prior to conducting an experiment safeguards against selective outcome reporting. This form of bias affects the quality of research in neurology. For example, when a randomized controlled trial (RCT) contains an outcome deemed “not significant” and is selectively removed from a trial, the validity of the RCT may be questioned. Previous studies have already established outcome reporting bias as an issue within neurology, noting that only 40% of analyzed RCTs were preregistered and, therefore, prespecified their analysis.[15] This same study found outcome reporting inconsistencies that often favored statistically significant results.[15] *JAMA Neurology, The Lancet Neurology*, and *Neurology* all requiring the preregistration of clinical trials prior to study commencement in accordance with the International Committee of Medical Journal Editors (ICJME).[39] Only *The Lancet Neurology* mentions registration of other study designs, such as observational studies, and only “encourages the registration of all observational studies on a WHO-compliant registry.”[40–42] The ICJME notes that although non-trial study designs lack a researcher prespecified intervention, it is recommended to preregister all study types to discourage selective reporting and selective publication of results [39]. On ClinicalTrials.gov alone, almost 65,000 observational study designs have been preregistered, comprising 21% of all registered studies [43]. Encouraging the preregistration of clinical trials and observational studies, alike, will increase transparency, increase the evidence available for systematic reviews and meta-analyses, and improve reproducibility [44, 45].

### Moving Forward

We propose the following solutions to promote reproducible and transparent research practices in neurology. With regards to journals, we recommend requiring open data sharing upon submission, or, at least, a statement from the authors signifying why open data sharing does not apply to their study. There are many open data repositories available, including the Open Science Framework (https://osf.io/), opendatarepository.org, and others listed at re3data.org. Second, we recommend journals and funding providers consider incentivizing reproducible research practices. For example, the Open Science Framework awards “badges” for open research practices, such as open data sharing, materials availability, and preregistration.[46] If one or more of these reproducible research practices do not apply to a particular study, a statement as to such should still qualify for the award. One Neuroscience journal, *Journal of Neurochemistry*, has already implemented open science badges with considerable success.[47] With regards to researchers, better awareness and education is necessary to encourage transparent and reproducible practices. Organizations, such as the *Global Biological Standards Institute*, have committed to improving the reproducibility of life sciences research through multiple methods, including training and educating researchers in effective trial design.[48, 49] The institute’s president has called for and implemented training programs aimed at teaching students, postdoctoral fellows, and principal investigators the importance of robust study design.[48] Additionally, we propose that medical schools and residency programs incorporate classes and didactic programs detailing proper experimental design with an emphasis on reproducible scientific practices. Research education should be a pillar of medical education, as physicians play an important role in guiding evidence-based healthcare. We anticipate that these recommendations, if implemented, will improve reproducibility within neurology and, as a result, the quality of research produced within this specialty.

### Strengths and Limitations

We feel that our methodology is robust and has many strengths, including blind and duplicate data extraction. Additionally, our protocol and data are available online to encourage reproducibility and transparency. However, we acknowledge a few limitations. First, we recognize that not all publications (clinical trials and protected patient data) are readily able to share their data and materials, although we feel a statement should still be reported. Second, we did not contact authors to obtain data, materials or analysis scripts and only used published materials for extraction. Had we contacted the authors, then source data, materials, and protocols may have been available.

## Conclusions

In summary, improvement is needed to incorporate reproducibility factors in neurology research. Such necessary improvement is attainable. Authors, journals, and peer-reviewers all have a part to play in developing an improved community of patient-centered neurology researchers. Reproducibility is paramount in evidence-based medicine to corroborate findings and ensure physicians have the highest quality evidence upon which to base patient care.

## List of Abbreviations

Randomized Control Trial (RCT)

## Declarations

Ethics Approval and Consent to Participate – Not Applicable

Consent for Publication – Not Applicable

Availability of Data and Materials – All protocols, materials, and raw data are available online via bioRxiv (BIOARKIV/2019/763730).

Competing Interests – We declare no conflicts of interest.

Funding – This study was funded through the 2019 Presidential Research Fellowship Mentor – Mentee Program at Oklahoma State University Center for Health Sciences.

Author’s Contributions – All authors read and approved the final manuscript. Individual roles detailed in methods section.

Acknowledgements – Not Applicable

Author’s Information – Not Applicable

---

**Supplemental 1:** Additional Characteristics of Reproducibility in Neurology Studies

| **Supplemental 1: Additional Characteristics of Reproducibility in Neurology** |  |  |  |
| --- | --- | --- | --- |
| **Studies** |  |  |  |
| **Characteristics** |  | **Variables** |  |
|  |  | **N (%)** | **95% CI** |
| **Conflict of Interest Statement (N=290)** | Statement, one or more conflicts of interest | 40 (13.79) | 9.89-17.70% |
|  | Statement, no conflict of interest | 154 (53.10) | 47.46-58.75% |
|  | No conflict of interest statement | 96 (33.10) | 27.78-38.43% |
| **Data Availability (N=202)** | Statement, some data are available | 19 (9.41) | 6.10-12.71% |
|  | Statement, data are not available | 0 | 0 |
|  | No data availability statement | 183 (90.59) | 87.29-93.90% |
| **Material Availability (N=189)** | Statement, some materials are available | 17 (8.99) | 5.76-12.23% |
|  | Statement, materials are not available | 0 | 0 |
|  | No materials availability statement | 172 (91.01) | 87.77-94.24% |
| **Protocol Availability (N=202)** | Full Protocol | 2 (0.99) | 0-2.11% |
|  | No Protocol | 200 (99.01) | 97.89-100% |
| **Analysis Scripts (N=202)** | Statement, some analysis scripts are available | 1 (0.50) | 0-1.29% |
|  | Statement, analysis scripts are not available | 0 | 0 |
|  | No analysis script availability statement | 201 (99.50) | 98.71-100% |
| **Replication**<br>**Studies**<br>**(N=202)** | Reports Replication Study | 0 | 0 |
|  | No Clear Statement | 202 (100) | 100% |
| **Open Access**<br>**(N=290)** | Yes - found via Open Access Button | 117 (40.34) | 34.79-45.90% |
|  | Yes - found article via other means | 5 (1.72) | 0.25-3.20% |
|  | Could not access through paywall | 178 (61.38) | 55.87-66.89% |
| **Cited in a**<br>**Systematic**<br>**Review/**<br>**Meta-Analysis (a)**<br>**(N=205)** | No Citations | 163 (79.51) | 74.94-84.08% |
|  | A Single Citation | 28 (13.66) | 9.77-17.54% |
|  | One to Five Citations | 13 (6.34) | 3.58-9.10% |
|  | More Than 5 Citations | 1 (0.49) | 0-1.28% |
| Abbreviations: CI, Confidence Interval. a - No studies were explicitly excluded from the systematic reviews or meta-analyses that cited the original article. |  |  |  |

**Supplemental 2:**
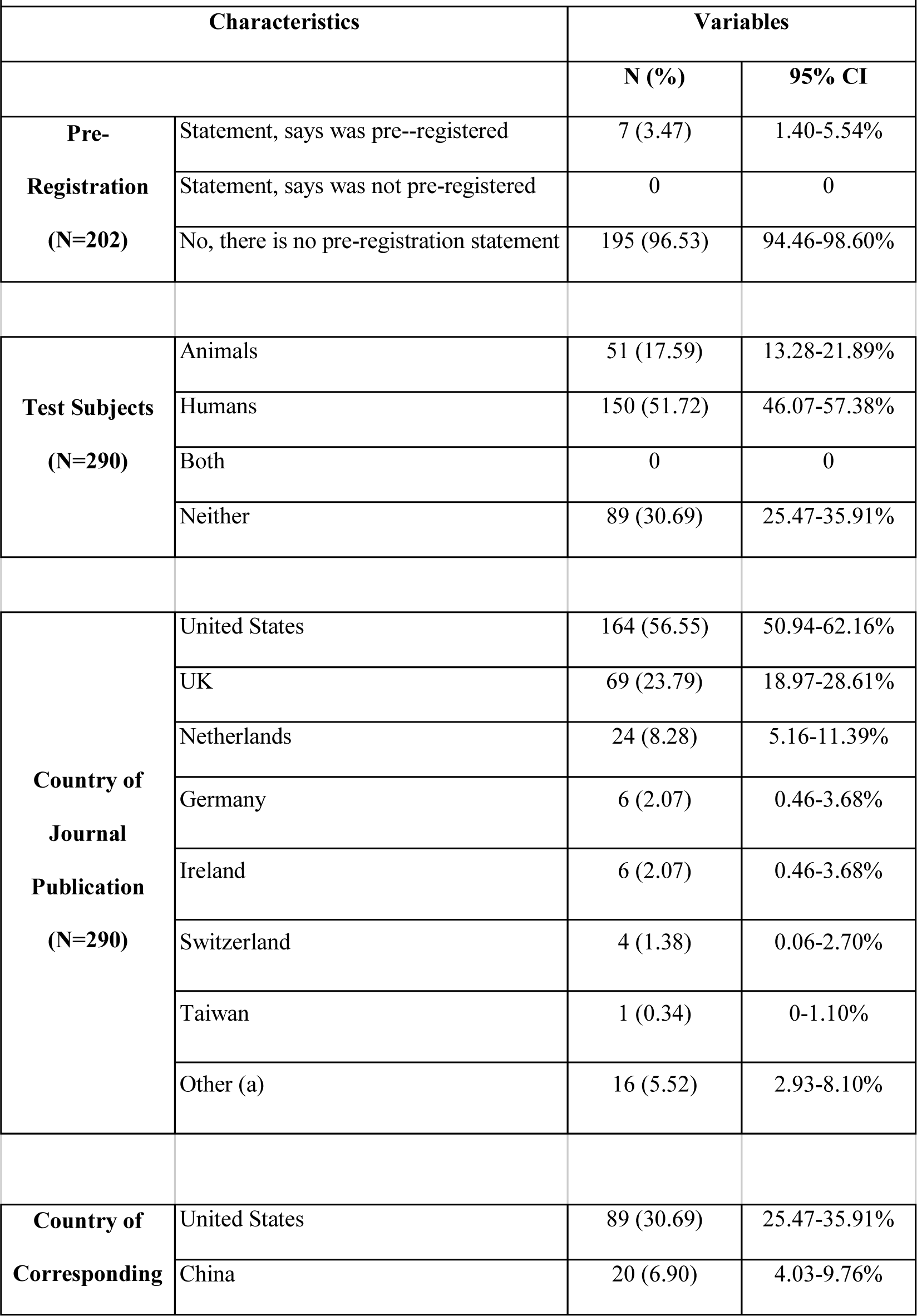
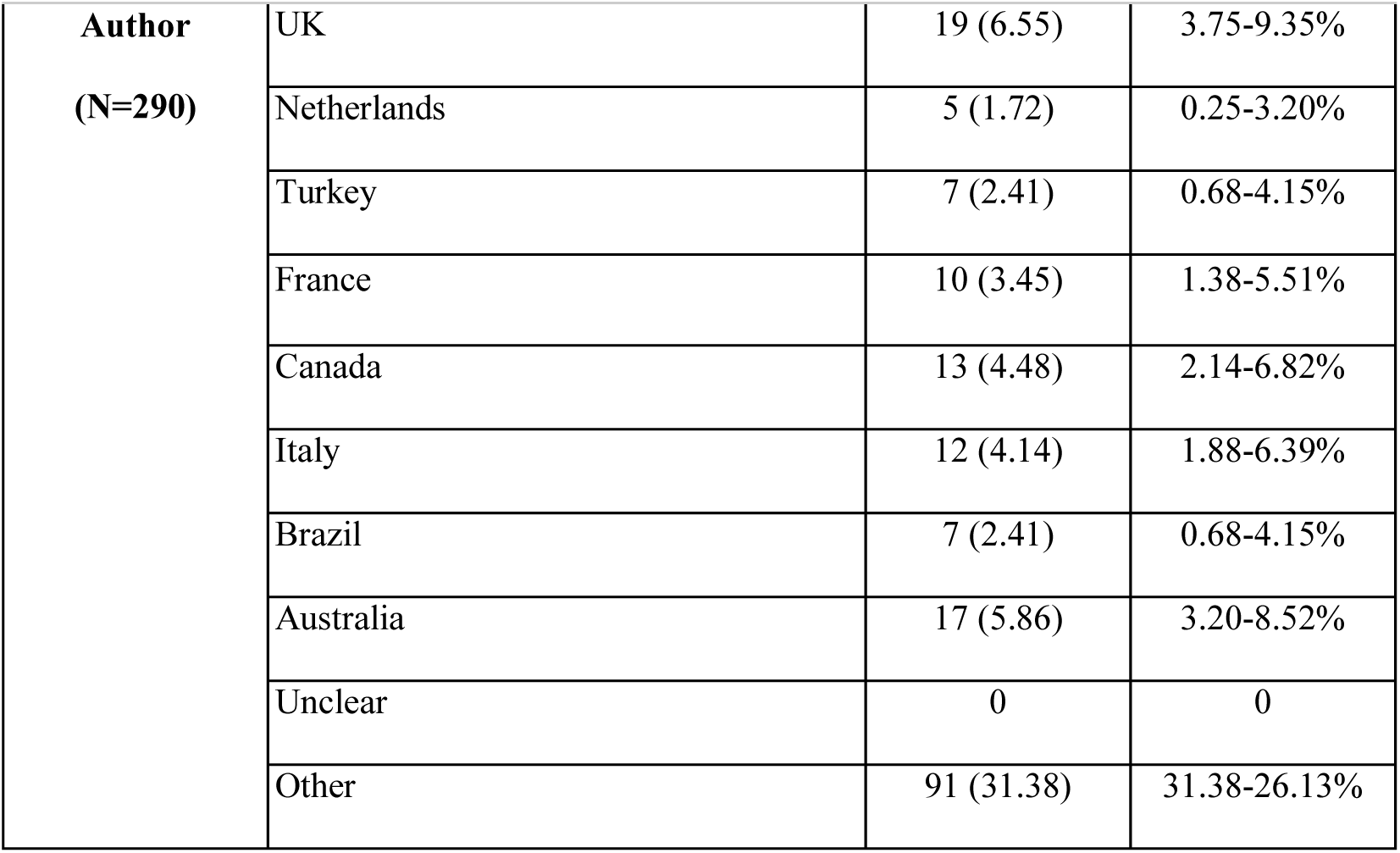
Additional Characteristics of Reproducibility in Neurology Studies

**Supplemental 3:** Additional Characteristics of Reproducibility in Neurology Studies

|  |  |  |
| --- | --- | --- |
| **Characteristics** |  | **Variables** |
|  |  | **N** |
| **Material**<br>**Availability** | Personal or institutional | 0 |
|  | Supplementary information hosted by the<br>journal | 13 |
|  | Online third party | 3 |
|  | Upon Request | 1 |
|  | Yes, material was accessible | 11 |
|  | No, material was not accessible | 6 |
| **Data**<br>**Availability** | Personal or institutional | 0 |
|  | Supplementary journal information | 11 |
|  | Online third party | 4 |
|  | Upon Request | 4 |
|  | Other (b) | 0 |
|  | Yes, data could be accessed and downloaded | 4 |
|  | No, data count not be accessed and downloaded | 15 |
|  | Yes, data files were clearly documented | 1 |
|  | No, data files were not clearly documented | 4 |
|  | Yes, data files contain all raw data | 1 |
|  | No, data files do not contain all raw data | 3 |
|  | Unclear if all raw data was available | 0 |
| **Pre-**<br>**Registration** | Yes, there was a pre-regisration | 7 |
|  | Registered on ClinicalTrials.Gov | 4 |
|  | Registered on Other | 3 |
|  | Hypothesis was pre-registered | 2 |
|  | Methods were pre-registered | 7 |
|  | Analysis plan was pre-registered | 6 |

